# The evolution of the gliotoxin biosynthetic gene cluster in *Penicillium* fungi

**DOI:** 10.1101/2023.01.17.524442

**Authors:** Charu Balamurugan, Jacob L. Steenwyk, Gustavo H. Goldman, Antonis Rokas

## Abstract

Fungi biosynthesize a diversity of secondary metabolites, small organic bioactive molecules that play diverse roles in fungal ecology. Fungal secondary metabolites are often encoded by physically clustered sets of genes known as biosynthetic gene clusters (BGCs). Fungi in the genus *Penicillium* produce diverse secondary metabolites that have been both useful (e.g., the antibiotic penicillin and the cholesterol-lowering drug mevastatin) and harmful (e.g., the mycotoxin patulin and the immunosuppressant gliotoxin) to human affairs. BGCs often also encode resistance genes that confer self-protection to the secondary metabolite-producing fungus. Some *Penicillium* species, such as *Penicillium lilacinoechinulatum* and *Penicillium decumbens*, are known to produce gliotoxin, a secondary metabolite with known immunosuppressant activity; however, an evolutionary characterization of the BGC responsible for gliotoxin biosynthesis among *Penicillium* species is lacking. Here, we examine the conservation of genes involved in gliotoxin biosynthesis and resistance in 35 *Penicillium* genomes from 23 species. We found homologous, less fragmented gliotoxin BGCs in 12 genomes, mostly fragmented remnants of the gliotoxin BGC in 21 genomes, whereas the remaining two *Penicillium* genomes lacked the gliotoxin BGC altogether. In contrast, we observed broad conservation of homologs of resistance genes that reside outside the BGC across *Penicillium* genomes. Evolutionary rate analysis revealed that BGCs with higher numbers of genes evolve slower than BGCs with few genes. Even though the gliotoxin BGC is fragmented to varying degrees in nearly all genomes examined, ancestral state reconstruction suggests that the ancestor of *Penicillium* species possessed the gliotoxin BGC. Our analyses suggest that genes that are part of BGCs can be retained in genomes long after the loss of secondary metabolite biosynthesis.

## Introduction

Gliotoxin is a secondary metabolite produced by certain fungi, including the major opportunistic human pathogen *Aspergillus fumigatus* (Raffa and Keller 2019). Secondary metabolites are bioactive molecules of low molecular weight that are not required for the organism’s growth but aid survival in harsh environments (Raffa and Keller 2019). Genes that participate in the biosynthesis of secondary metabolites, including gliotoxin, typically reside next to each other in fungal genomes and form biosynthetic gene clusters (BGCs) (Rokas *et al*. 2020). The gliotoxin BGC is implicated in human pathogenicity because gliotoxin suppresses the immune response of the mammalian host through diverse mechanisms, including by inhibiting protein complexes necessary for the generation of antimicrobial reactive oxygen species, decreasing cytotoxic activities of T lymphocytes, and preventing integrin activation (Dolan *et al*. 2015; Raffa and Keller 2019). Gliotoxin’s role in modulating host biology suggests that it is a virulence factor (Raffa and Keller 2019). For example, virulence is attenuated in certain animal models of disease when *gliP*, the non-ribosomal peptide synthetase gene involved in gliotoxin biosynthesis, is deleted (Sugui *et al*. 2007).

Fungi that produce gliotoxin need to be resistant to the toxin. Several genes contribute to resistance, such as the thioredoxin reductase gene *gliT*, located within the gliotoxin BGC (Schrettl *et al*. 2010). *gliT* deletion strains of *A. fumigatus* exhibit resistance to gliotoxin oxidation and unchecked methylation (Owens *et al*. 2015). As a result, *gliT*-deficient *A. fumigatus* are hypersensitive to gliotoxin (Owens *et al*. 2015). Other resistance genes encoding transcription factors, transporters, and oxidoreductases, reside outside the BGC and – like *gliT* – are found in both gliotoxin-producing and non-producing species (Castro *et al*. 2022). For example, the transcription factor RglT is the primary regulator of *gliT* (Ries *et al*. 2020). Seven other genes are known to be regulated by *rglT* and contribute to gliotoxin resistance: *gtmA* (encodes a *bis*-thiomethyltransferase, AFUA_2G11120), *kojR* (transcription factor, AFUA_5G06800), *abcC1* (ABC-transporter, AN7879/AFUA_1G10390), *mtrA* (methyltransferase, AN3717/AFUA_6G12780), AN9051 (oxidoreductase, AFUA_7G00700), AN1472 (MFS transporter, AFUA_8G04630), and AN9531 (NmrA-transcription factor, AFUA_7G06920) (Castro *et al*. 2022).

Though progress has been made in understanding the mechanisms and functions of the gliotoxin biosynthetic pathway, several questions remain, especially concerning the evolutionary and ecological significance of this BGC in lineages that contain a mix of biotechnologically and medically relevant fungi, such as *Penicillium* (Steenwyk *et al*. 2019). For example, *Penicillium camemberti* and *Penicillium roqueforti* contribute to cheese production (Nelson 1970; Lessard *et al*. 2012), whereas *Penicillium expansum*, *Penicillium digitatum*, and *Penicillium italicum* are postharvest pathogens of citrus fruits, stored grains, and other cereal crops (Marcet-Houben *et al*. 2012; Ballester *et al*. 2015; Li *et al*. 2015). Examination of the gliotoxin BGC in the genomes of *Penicillium* species will shed light on the evolution of the gliotoxin BGC within Aspergillaceae, the family encompassing both *Aspergillus* and *Penicillium* species.

Considering the close relatedness of *Penicillium* and *Aspergillus*, it is interesting that evidence of gliotoxin production is scant within the former. To fill this gap, we employed a genome-scale approach to infer the evolutionary history of the gliotoxin BGC among 35 strains of 23 *Penicillium* species. We found that most *Penicillium* genomes examined contained fragmented gliotoxin BGCs and two lacked a BGC. However, some *P. expansum* strains had two homologous gliotoxin BGCs. Codon optimization analysis reveals that genes *in Penicillium* BGCs are lowly optimized, whereas genes in *Aspergillus* gliotoxin BGCs are highly optimized.

In contrast, gliotoxin resistance genes in *Penicillium* and *Aspergillus* fungi have similar degrees of codon optimization, suggesting that *Penicillium* species encounter exogenous gliotoxin in their environments. Examination of evolutionary rates revealed that genes from highly fragmented gliotoxin BGCs evolved at significantly higher rates than genes from less fragmented BGCs, suggesting that less fragmented BGCs have been experiencing relaxation of selective constraints for longer. Ancestral state reconstructions indicate that the *Penicillium* ancestor possessed a less fragmented gliotoxin BGC, followed by distinct trajectories of duplication and loss, highlighting the diverse evolutionary pathways of the gliotoxin BGC in *Penicillium* species.

## Materials and Methods

### I. Data collection and quality assessment

We retrieved the genomes and gene annotations of 35 *Penicillium* strains from 23 species as well as of two outgroups (*Aspergillus fumigatus* and *Aspergillus fischeri*) from NCBI (https://www.ncbi.nlm.nih.gov/) (Table S1).

Genome assembly and annotation quality were examined to evaluate whether the dataset is sufficient for comparative genomics. The quality and characteristics of the genomes (N50, L50, assembly size, number of scaffolds, and gene count) were evaluated using BioKIT (v0.1.0) (Steenwyk *et al*. 2022) (Figure S1). The average N50 value was 1,850,972.1 bases, where 46% of proteomes consisted of N50 values greater than 1 Mb, and the lowest N50 value was 31,119 bases for *P. expansum* CMP 1. Gene annotation completeness was assessed using BUSCO (v5.0.0) (Waterhouse *et al*. 2018) (Figure S2). BUSCO uses a predetermined set of near-universally conserved single-copy genes (or BUSCO genes) to identify their presence in a query proteome (characterized as single-copy, duplicated, or fragmented) or absence. We used the 4,181 BUSCO genes from the Eurotiales OrthoDB dataset (Manni *et al*. 2021; Zdobnov *et al*.2021). Nearly all the genomes have high BUSCO gene coverage (average: 95.9% ± 3.1%), with the lowest percentages being for *P. coprophilum* (87.9%) and *P. decumbens* (85.3%).

### II. Identification and characterization of gliotoxin BGC and resistance genes

#### a. Identification of gliotoxin GBC and resistance genes

The representative gliotoxin BGC (BGC0000361, Download date: April 2022) from the *Aspergillus fumigatus* Af293 reference strain was downloaded from the Minimum Information about a Biosynthetic Gene Cluster (MiBIG) database (Kautsar *et al*. 2019). Command-line NCBI BLASTP (Camacho *et al*. 2009) searches for the Af293 gliotoxin BGC against the proteome of each species were executed. Highly similar sequences were identified using an expectation value threshold of 1e-4 and a query coverage of 50%. The resulting BLAST outputs were then cross-referenced with the NCBI feature table file, which contains genome location information for each gene, and parsed to identify clusters of homologs. Less fragmented BGCs are defined as having at least 7 / 13 genes from the query gliotoxin BGC present, including *gliP*, encoding the core nonribosomal peptide synthetase (Castro *et al*. 2022); mostly fragmented clusters are defined as having at least 3 /13 genes from the gliotoxin BGC without a requirement for this cluster to include *gliP*. When identifying BGCs, we allowed up to four genes between each pair of adjacent homologs using the *A. fumigatus* Af293 BGC from the MiBIG database (Kautsar *et al*. 2019) as reference (Castro *et al*. 2022).

To rule out gene annotation errors in cases where we infer genes to be absent, we conducted command-line NCBI tBLASTn searches for the Af293 gliotoxin BGC against the genome sequences. Highly similar sequences were identified using an expectation value threshold of 1e-10. The resulting outputs were analyzed, and no new presence/absence information was found.

Sequence similarity searches were also conducted for eight gliotoxin resistance genes (*abcC1*/AN7879, *mtrA/* AN3717, AN9051, AN1472, AN9531, *rglT, gtmA*/AFU2G11120, *kojR*/AFUA_5G06800), three of which were transcription factors (AN9531, *rglT, kojR*). We used an expectation value threshold of 1e-3 and a query coverage threshold of 50%; we used a lower query coverage threshold of 40% for the three transcription factors.

#### b. Codon bias

To estimate the potential functional significance of the partial gliotoxin BGCs present in *Penicillium* genomes, mean gene-wise relative synonymous codon usage (gRSCU) was determined for each clustered *gli* gene across all proteomes using BioKIT (Steenwyk *et al*.2022). This provides insight into how codon usage bias influences the expression level of a particular gene. The percentile rankings of each of the present and clustered *gli* genes were calculated using the R package *dplyr* (v1.0.9) (Wickham *et al*. 2022), and these values, for each species, were then plotted using the R package *ggplot2* (Wickham 2016).

#### c. Synteny Analysis

Alignments of representative *Penicillium* genomes with less and more fragmented gliotoxin BGCs were generated using a GenomeDiagram in Biopython (Cock *et al*. 2009). Five genomes (*A. fumigatus* Af293, *P. flavigenum* IBT 14082, *P. roqueforti* FM164, *P. nordicum* DAOMC 185683, and *P. expansum* CMP1) with the largest number of different, homologous *gli* cluster genes above seven, and including *gliP*, were chosen to visualize the conservation of synteny of less fragmented gliotoxin BGCs across the phylogeny. Similarly, the five genomes (*P. steckii* IBT 24891, *P. vulpinum* IBT 29486, *P. rubens* 43M1, *P. camemberti* FM 013, and *P. italicum* PHI 1) with the greatest number of different, homologous *gli* cluster genes above three and below seven, and not needing to include *gliP*, were chosen to visualize synteny of mostly fragmented BGCs across the phylogeny.

### III. Phylogenetic Analysis

#### a. Species Tree Inference

The evolutionary relationships of *Penicillium* species were obtained from a previous study (Steenwyk *et al*. 2019) using treehouse (Steenwyk and Rokas 2019). For three species with population-level data, within-species relationships were inferred using phylogenomics. To do so, protein sequences of BUSCO genes were first aligned using MAFFT (v7.490) with the *--auto* parameter (Katoh and Standley 2013). Codon-based alignments were generated by threading the corresponding DNA sequences onto the protein alignment with the *thread_dna* function in PhyKIT (v1.11.2) (Steenwyk *et al*. 2021). The resulting nucleotide alignments were trimmed using ClipKIT (v1.3.0) (Steenwyk *et al*. 2020) with default parameters. The resulting aligned and trimmed sequences were concatenated into a supermatrix with 8,124,861 sites using the *create_concat* function in PhyKIT. We then inputted the concatenated matrix into IQ-TREE 2 (v2.0.6), a software that implements a maximum likelihood framework for inferring phylogenies. All other evolutionary relationships between species were constrained following the relationships inferred in a previously published study (Steenwyk and Rokas 2019). The best-fitting substitution model (GTR+F+I+G4) was determined using ModelFinder (Kalyaanamoorthy *et al*.2017).

#### b. Single-gene tree inference

To infer the evolutionary history of genes in the gliotoxin BGCs, individual *gli* genes were compiled and aligned with MAFFT (v7.490) using the *--auto* parameter (Katoh and Standley 2013). The corresponding nucleotide sequences for each file were obtained from the CDS files for each species, using the *faidx* function of BioKIT (v0.1.0) (Steenwyk *et al*. 2022). These nucleotide sequences were then threaded onto the protein alignments using the *thread_dna* function of PhyKIT (Steenwyk *et al*. 2021), resulting in a codon-based alignment. All individual codon-based gene alignments were trimmed with ClipKIT (Steenwyk *et al*. 2020) with default parameters. The trimmed alignments were used to construct a phylogeny using IQ-TREE 2 (Minh *et al*. 2020). The best-fitting substitution model was chosen for each *gli* gene using Bayesian information criteria (BIC) implemented in ModelFinder (Kalyaanamoorthy *et al*. 2017) from IQ-TREE 2. Branch support in each phylogenetic tree was assessed by 1000 bootstraps using ultrafast bootstrapping approximation (Hoang *et al*. 2018). Tree visualization was carried out using the R packages *ape* (v5.6.2) (Paradis and Schliep 2019) and *phytools* (v1.0.3) (Revell 2012).

To characterize variation in the evolution of individual genes of the gliotoxin BGC, the trimmed alignments and maximum-likelihood trees from IQ-TREE 2 were used as input into the *evolutionary_rate, total tree length*, and *pairwise identity* functions of PhyKIT to estimate two tree-based measures of evolutionary rate and one sequence-based measure. Evolutionary rate is defined as the total tree length divided by the number of terminals (Telford *et al*. 2014; Steenwyk *et al*. 2021). The total tree length is the sum of all branches (Steenwyk *et al*. 2021).

#### c. Ancestral state reconstructions

Ancestral state reconstruction for each gene of the gliotoxin BGC across three discrete characters (“Presence clustered,” “Presence unclustered,” and “Absence”) was estimated using *phytools* (v1.0.3) (Revell 2012). Presence generally indicates that a homolog of the particular gene was identified. “Presence clustered” identifies an existing homolog of the specific gene within a maximum distance of four genes from other homologs of the gliotoxin BGC. “Presence unclustered” identifies an existing homolog of the particular gene without clustering. “Absence” indicates that no homolog of the specific gene was identified. Estimation of ancestral character states was done using the Dollo parsimony method. This method assumes that a complex character lost during the evolution of a particular lineage cannot be regained (Rogozin *et al*.2006). Count, a software package for the evolutionary analysis of homolog family sizes, was used to generate these ancestral state reconstructions (Csűös 2010).

#### d. Tree Topology Testing

Tree topology testing was used to determine whether the duplication event resulting in the two less fragmented, homologous gliotoxin BGCs in *P. expansum* strains MD 8 and d1 occurred solely in the lineage of *P. expansum* or deeper in the tree. IQTREE 2 (Minh *et al*. 2020) was used to compute log-likelihoods of a constrained tree (monophyly of *P. expansum gliP* homologs) and the observed tree in which a polyphyly of *gliP* in both clusters is seen (inconsistent with the known species tree). 1000 RELL replicates (Kishino *et al*. 1990) were performed. The AU test results (Shimodaira 2002) was used for comparison.

## Results and Discussion

### I. The gliotoxin BGC is fragmented in *Penicillium* species

Presence / absence data of the 13 genes in the gliotoxin BGC among the 23 *Penicillium* species analyzed reveals that the cluster is largely fragmented in the genus *Penicillium* (Figure 1). The proteomes of 12 strains from 5 *Penicillium* species (*P. arizonense, P. flavigenum, P. roqueforti, P. nordicum, P. expansum*), possessed less fragmented BGCs, and the proteomes of 23 strains from 18 *Penicillium* species had mostly fragmented BGCs (Figure S3-S15). Two less fragmented BGCs, which contained 10 / 13 genes and 7 / 13 genes, were identified in *P. expansum* strains d1 and MD 8, respectively. Regardless of the number of less fragmented BGCs found, to our knowledge, none of the *Penicillium* species in question are known to produce gliotoxin, except *P. decumbens* (Feng *et al*. 2018), suggesting that the absence of clustering in this species may be due to strain heterogeneity and requires further exploration.

**Fig. 1.**
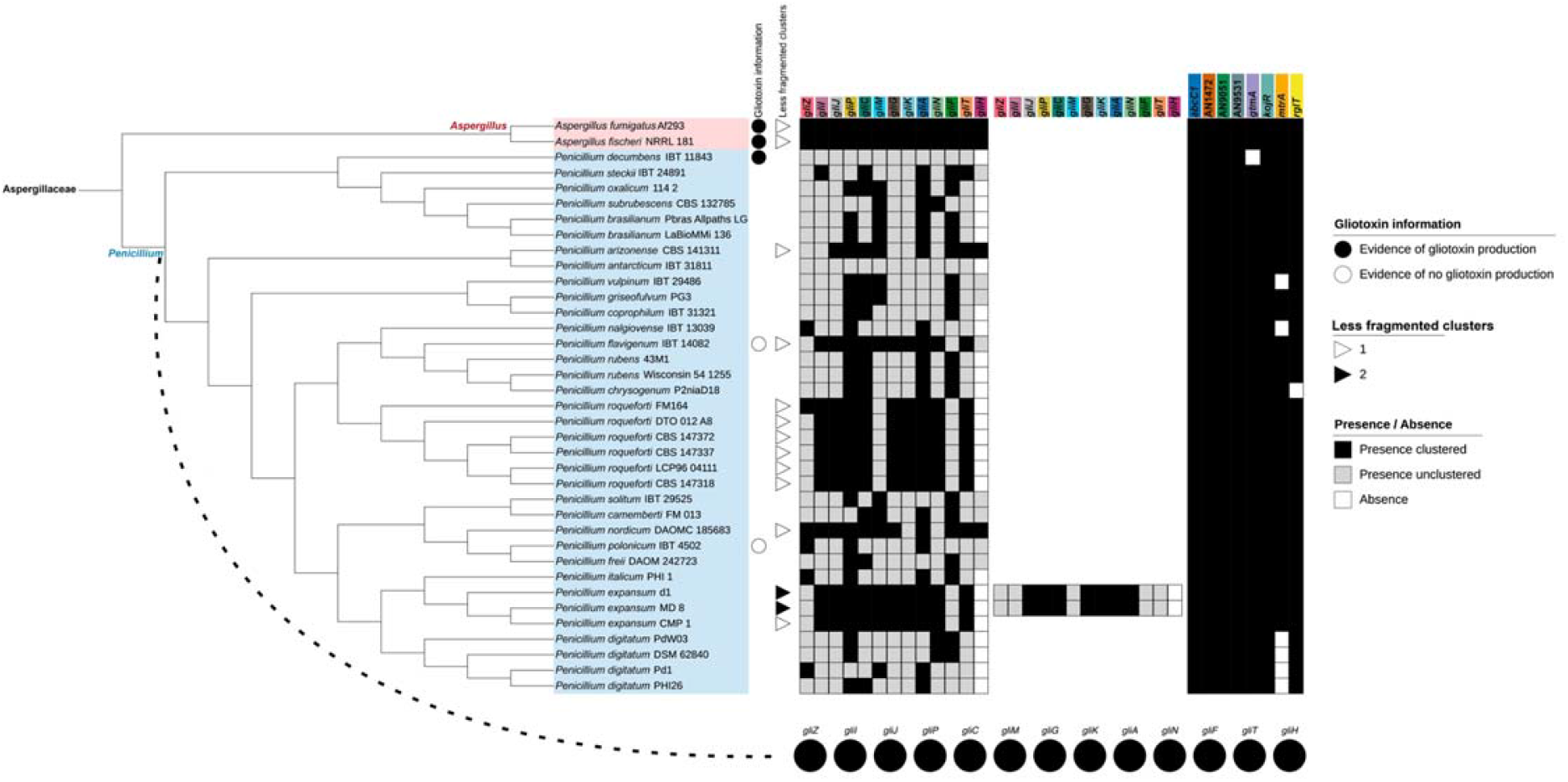
Phylogeny of *Penicillium* genomes. Different genera are depicted using different-colored boxes. *Aspergillus* is shown in red and *Penicillium* in blue. Shaded circles next to species / strain names indicate gliotoxin production information from the literature, or lack thereof (Fischer *et al*. 2000; Spikes *et al*. 2008; Knowles *et al*. 2020; Redrado *et al*. 2022). Shaded squares in the second column depict number of clusters identified. Remaining color strips depict gene presence clustered (black), presence unclustered (gray), and absence (white) according to the requirements outlined in the *Methods* section. Ancestral state reconstructions of each gene of the gliotoxin BGC (for the ancestor of *Penicillium* species) are presented in pie charts below the phylogeny.

### II. A complete gliotoxin BGC was present in the ancestor of *Penicillium* species

Ancestral state reconstruction revealed the presence of all 13 genes in the gliotoxin BGC in the ancestor of the *Penicillium* species used in our study (Figure 1). We infer that the first gene lost was *gliH*, which is absent from 25 of the 35 *Penicillium* strains examined. The *gliH* gene encodes an acetyltransferase that, when deleted, results in a loss of gliotoxin production in *A. fumigatus* (Schrettl *et al*. 2010; Castro *et al*. 2022). Thus, the early loss of *gliH* in the genus *Penicillium* may have been the key determinant of a lack of gliotoxin production. Further, the synteny of genes in the BGC is mostly conserved and similar to the arrangement of the *A. fumigatus* Af293 gliotoxin BGC across representative, less fragmented BGCs, such as *P. flavigenum* IBT 14082 and *P. expansum* CMP 1 (Figure 2). In contrast, there is extensive divergence in synteny conservation among mostly fragmented BGCs (Figure 2). To our knowledge, none of the *Penicillium* species examined are known to produce gliotoxin, except *P. decumbens* (Feng *et al*. 2018), suggesting that the absence of clustering in this species may be due to strain heterogeneity and requires further exploration.

**Fig. 2.**
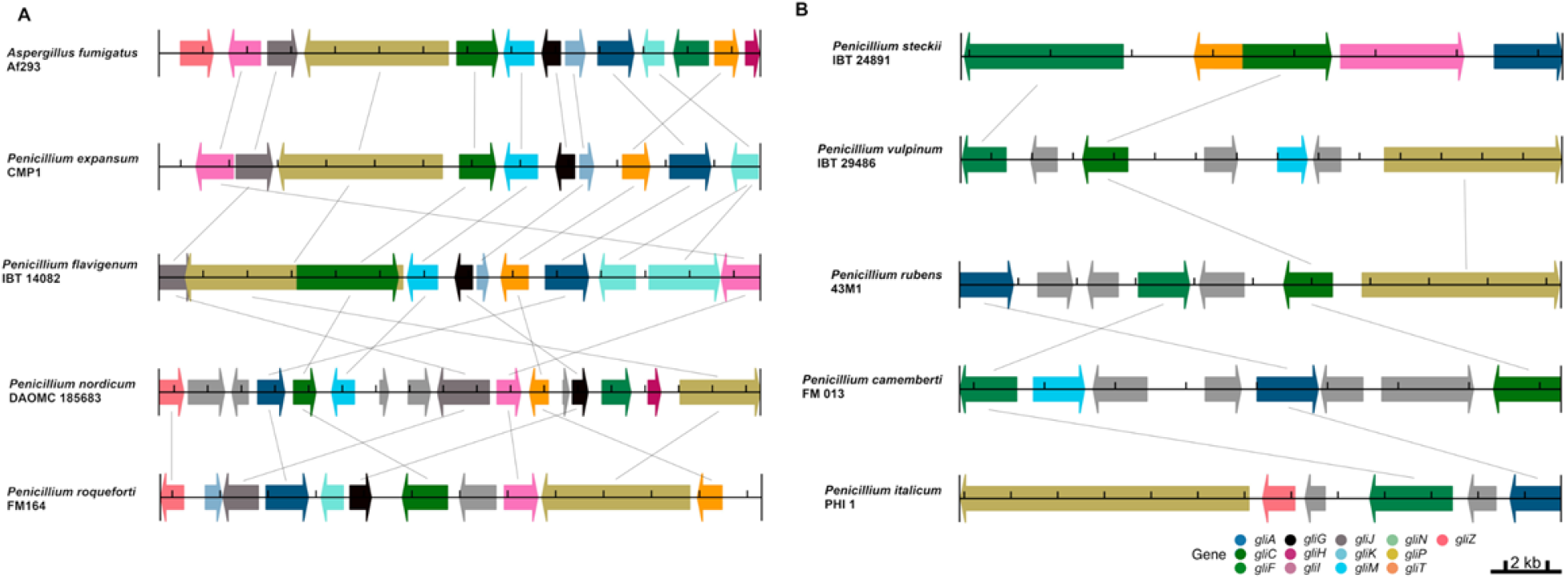
Conservation of gliotoxin BGC synteny for representative *Penicillium* species. Synteny analysis of representative genomes with less fragmented **(A)** and mostly fragmented **(B)** gliotoxin BGCs. Each interval along the track represents 2 kb.

### III. Resistance genes are broadly conserved

The presence/absence results of the eight resistance genes, portrayed in Figure 1, suggest that their origins predate the *Aspergillus* and *Penicillium* genera (Figure S16-S23). All species possessed *abcC1*, AN1472, AN9051, AN9531, and *kojR* homologs. In addition, only *Penicillium* species with mostly fragmented gliotoxin BGCs lacked at least one resistance gene, such as *gtmA*, *mtrA*, and *rglT*. *Penicillium chrysogenum* lacked both *rglT* and *gliT*, an observation consistent with the transcriptional dependency of *gliT* to *rglT* (Ries *et al*. 2020).

### IV. *Penicillium* species have experienced changes in gliotoxin BGC synteny over time

All genes of the gliotoxin BGC were broadly found within the genus *Penicillium*, except for *gliH*, yet most were sparsely clustered (Figure 1). More specifically, 12 out of 35 *Penicillium* species/strains were found to have a less fragmented, homologous BGC. Two strains of *Penicillium expansum* (d1 and MD 8) were found to have two BGCs. Evidence of variation in gene presence / absence is also evident within species. For example, *Penicillium roqueforti* shows population variation in the presence of *gliZ*, a major transcriptional regulator of gliotoxin biosynthesis (Bok *et al*. 2006); five strains of *P. roqueforti* lack *gliZ* whereas one strain has the gene. As a result, we can conclude that the ancestor of *P. roqueforti* had a *gliZ* homolog, but the gene was lost over time in most of the strains, highlighting the importance of population-level sampling. Overall, we can see that the gliotoxin BGC has experienced relocations and duplications of its genes, specifically in *Penicillium expansum* strains d1 and MD 8, as is expected in the formation of most secondary metabolite-producing BGCs (Rokas *et al*. 2018).

### V. Few *Penicillium* species contain codon-optimized gliotoxin BGCs

Compared to the two outgroup *Aspergillus* species, *A. fumigatus* and *A. fischeri*, *Penicillium* species have much lower gRSCU value rankings (Figure 3). Specifically, the mean gRSCU percentile rank of gliotoxin BGC genes among the *Aspergillus* outgroups is 0.81, while that among the *Penicillium* species is 0.35; these scores suggest that *gli* genes from *Aspergillus* are more codon-optimized than *gli* genes from *Penicillium*. Regardless of mean gRSCU values, *gliT* and *gliA* homologs, when present, are ranked consistently in the top three to four clustered genes. However, when considering resistance genes, the spread and range of their gRSCU values are similar across all species. The mean gRSCU percentile rank of gliotoxin resistance genes among the *Aspergillus* outgroups is 0.58, while that among the *Penicillium* species is 0.53. This allows us to infer that these *Penicillium* species may ecologically encounter exogenous gliotoxin, making *gliT*, encoding a gliotoxin-neutralizing enzyme, *gliA*, encoding a transporter that exports gliotoxin, and non-TF resistance genes such as *abcC1*, encoding an ABC-transporter, rank in the top percentiles among each of the species’ gene sets.

**Fig. 3.**
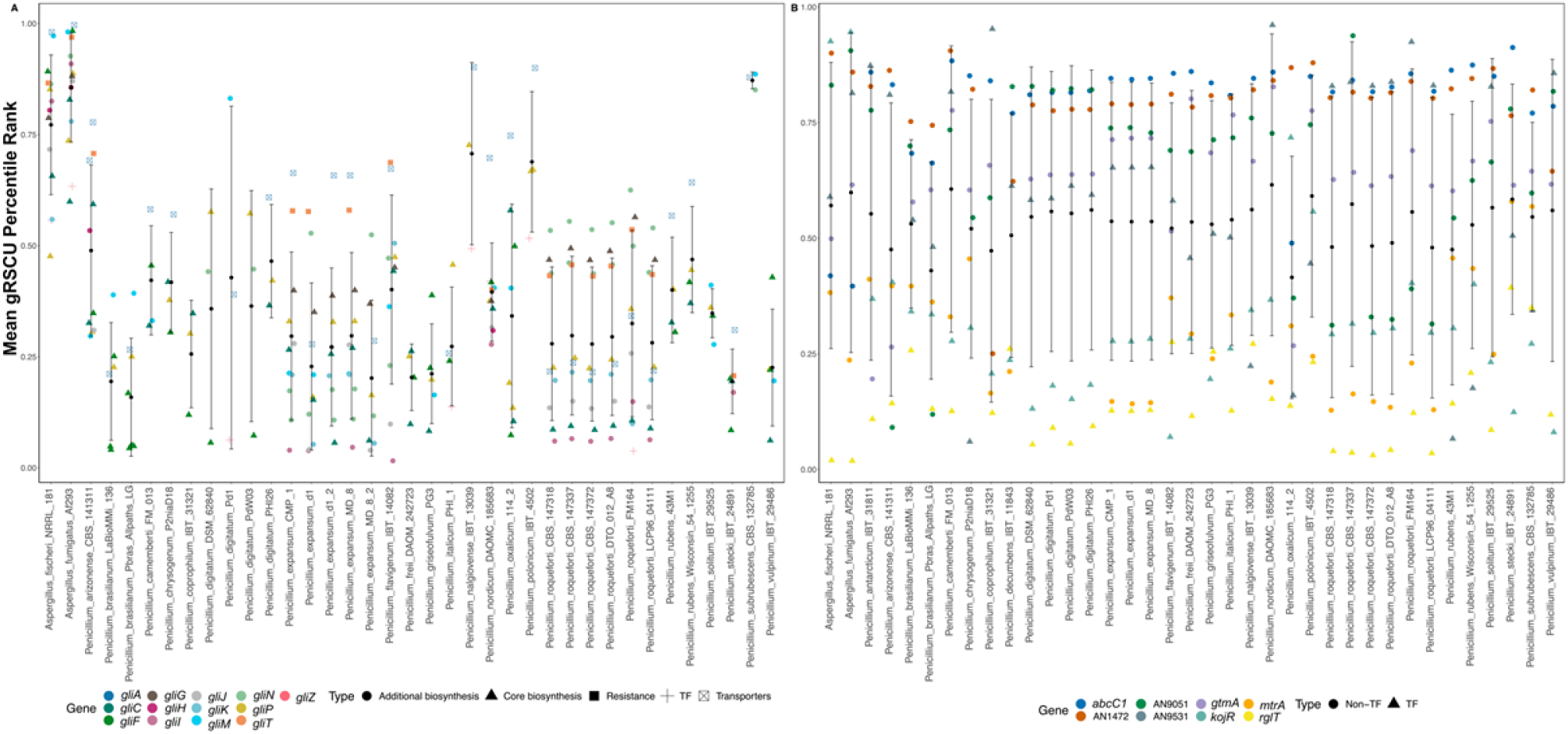
Gene-wise relative synonymous codon usage (gRSCU) for gliotoxin BGC and resistance genes. **(A)** Percentile rankings of gene-wise relative synonymous codon usage (gRSCU) among gliotoxin BGC genes, in comparison to all other genes. Types / functionality of each gene of the gliotoxin BGC is depicted by shape in the categories of “Core biosynthesis”, “Additional biosynthesis”, “Resistance”, “Transcription Factor”, “Transporter” **(B)** Percentile ranking of gene-wise relative synonymous codon usage (gRSCU) among gliotoxin resistance genes, in comparison to all other genes. Types / functionality of each resistance gene is depicted by shape in the categories of “Non-Transcription Factor and Transcription Factor”.

### VI. *gli* genes in less fragmented clusters are evolving at a slower rate than mostly fragmented clusters

In the comparison of tree-based and sequence-based measures of evolutionary rate, *gli* genes from less fragmented clusters are evolving at a significantly slower pace (p<0.0001) than those from mostly fragmented clusters across all three metrics, as seen by a two-way ANOVA with an additive model (Figure 4, Figure S3-S15). This difference highlights a notable feature of many BGCs, the fact that they are rapidly evolving, hinted at by their high variability and narrow taxonomic range (Rokas *et al*. 2020).

**Fig. 4.**
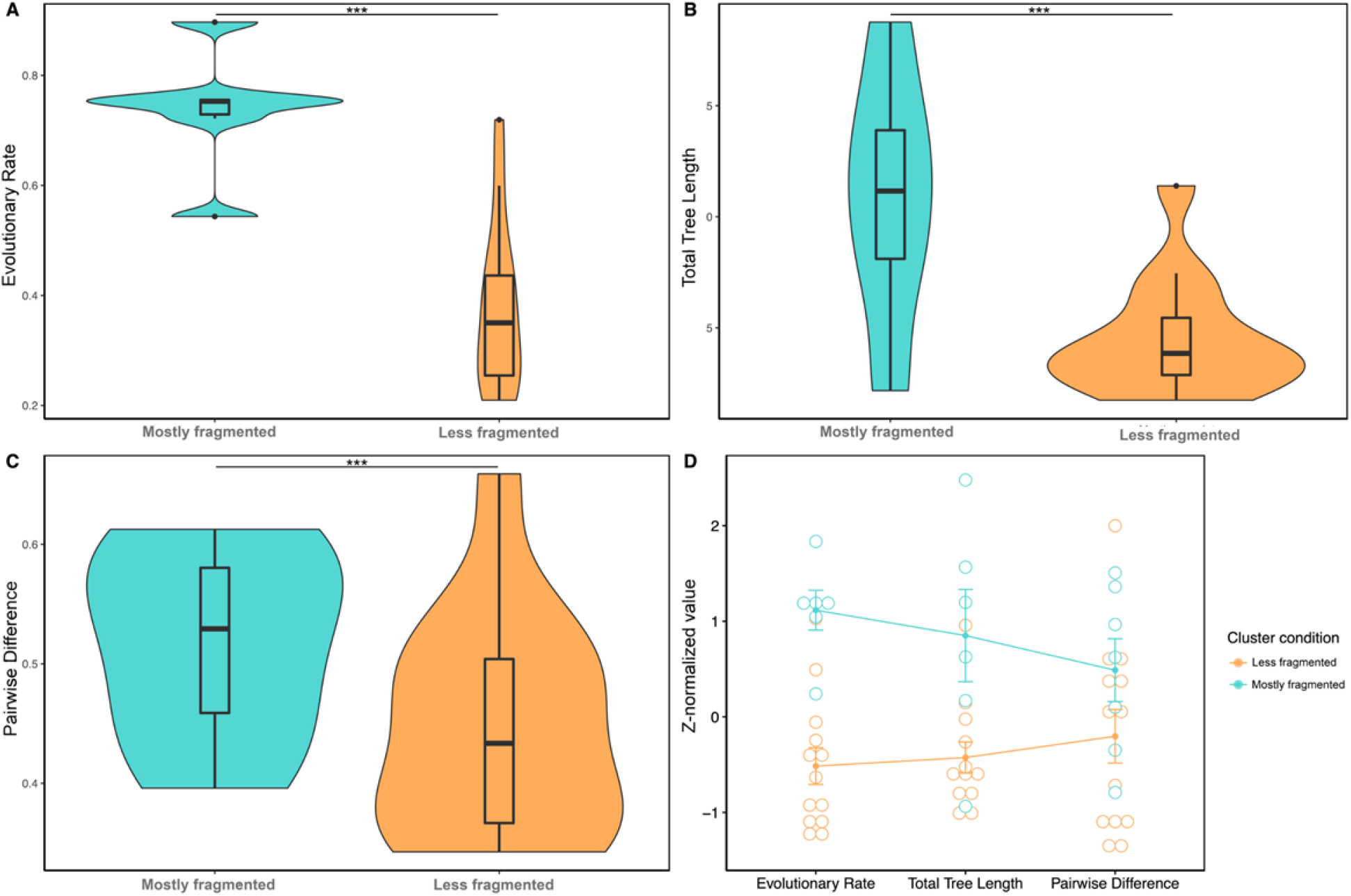
Evolutionary rate comparison across gliotoxin BGCs. Multi-method comparison of evolutionary rates between less fragmented and mostly fragmented gliotoxin BGCs. Less fragmented clusters were required to contain a *gliP* ortholog and at least 7 different genes of the cluster. Mostly fragmented clusters had no requirement to contain a *gliP* ortholog and only needed to contain at least 3 different genes of the cluster. **(A)** Comparison of evolutionary rates, as a function of total tree length divided by the number of taxa, between less fragmented and mostly fragmented gliotoxin BGCs. **(B)** Comparison of total tree length between less fragmented and mostly fragmented gliotoxin BGCs. **(C)** Comparison of pairwise identity between less fragmented and mostly fragmented gliotoxin BGCs.

### VII A duplication of the gliotoxin BGC may have occurred before the divergence between *P. flavigenum* and *P. roqueforti*

We conducted a tree topology test to infer when the gliotoxin BGCs found in *P. flavigenum* occurred. The maximum likelihood phylogeny suggests that this duplication occurred before the divergence between *P. flavigenum* and *P. roqueforti*. An alternative hypothesis is that duplication occurred within *P. expansum*. This alternative hypothesis would be supported by monophyly of *P. expansum* homologs of BGC genes. After conducting a tree topology test comparing log likelihood values between the maximum likelihood phylogeny and an alternative tree wherein *P. expansum* gliP homologs were constrained to be monophyletic, we found that the constrained topology was significantly rejected (Approximately Unbiased test, p = 7.34e-110) (Figure S24). In other words, it is unlikely duplication occurred within *P. expansum* lineage; instead, duplication likely occurred more anciently, prior to the diversification of *P. expansum*.

## Conclusions

The ancestor of *Penicillium* species likely possessed a complete gliotoxin BGC. A duplication event of the BGC occurred in one lineage, likely prior to the divergence of *P. flavigenum* and *P. roqueforti*. Also, the presence/absence results of the eight resistance genes suggest that their origins predate the *Aspergillus* and *Penicillium* genera suggesting that resistance has long been important among these species. The genes in *Penicillium* gliotoxin BGCs are less codon optimized (gRSCU percentile rank mean: 0.35) compared to their *Aspergillus* counterparts (gRSCU percentile rank mean: 0.81) suggesting that *gli* genes are much more often expressed in *Aspergillus* species than in *Penicillium*. However, less fragmented BGCs within *Penicillium* species are evolving at a slower rate than mostly fragmented clusters, suggestive of potential functionality.

Although informative, this work only utilizes publicly available protein annotations of biotechnologically and medically relevant *Penicillium* fungi, making it important to expand upon the species/strains studied. Moreover, this same targeted gliotoxin analysis within a larger phylogeny of *Aspergillus* species, for which there is greater evidence of the production of this secondary metabolite, may be helpful. An analysis of gliotoxin BGCs encoded in all fungi would also provide us with more insight into the evolutionary mechanisms that give rise to BGC diversity. In addition, expanding on the causes of conservation of less fragmented gliotoxin BGCs within a variety of *Penicillium* strains may be important, especially because evidence of production is lacking. As a result, this exciting reality encourages further understanding of the motivating hypothesis that individual secondary metabolites are “cards” of virulence in a larger “hand” that fungi possess.

## Data availability

The authors affirm that all data necessary for confirming the conclusions of the article are present within the article, figures, tables, and supplemental material.

## Acknowledgements

We thank members of the Rokas Laboratory at Vanderbilt University for support and feedback on this work. We also thank the Vanderbilt Data Science Institute for their undergraduate enrichment opportunities. This work was performed in part using resources contained within the Advanced Computing Center for research and Education at Vanderbilt University in Nashville, TN.

## Funding

C.B. was supported by the Vanderbilt University Data Science Institute—Summer Research Program. J.L.S. and A.R. were funded by the Howard Hughes Medical Institute through the James H. Gilliam Fellowships for Advanced Study program. Research in A.R.’s lab is supported by grants from the National Science Foundation (DEB-2110404), the National Institutes of Health/National Institute of Allergy and Infectious Diseases (R01 AI153356), and the Burroughs Wellcome Fund.

## Conflicts of interest

J.L.S. is a scientific consultant for Latch AI Inc. J.L.S. is a scientific advisor for WittGen Biotechnologies. A.R. is a scientific consultant for LifeMine Therapeutics, Inc.

